# Diffusion tensor imaging and arterial tissue: establishing the influence of arterial tissue microstructure on fractional anisotropy, mean diffusivity and tractography

**DOI:** 10.1101/2020.05.20.104711

**Authors:** B. Tornifoglio, A. J. Stone, R. D. Johnston, S. S. Shahid, C. Kerskens, C. Lally

**Affiliations:** Trinity Centre for Biomedical Engineering, Trinity Biomedical Sciences Institute, Trinity College Dublin, Dublin, Ireland; Department of Mechanical and Manufacturing Engineering, School of Engineering, Trinity College Dublin, Dublin, Ireland; Department of Radiology and Imaging Sciences, Indiana University School of Medicine, Indianapolis, Indiana, USA; Trinity College Institute of Neuroscience, Trinity College Dublin, Dublin, Ireland; Advanced Materials and Bioengineering Research Centre (AMBER), Royal College of Surgeons in Ireland and Trinity College Dublin, Dublin, Ireland

**Keywords:** diffusion tensor imaging, arterial microstructure, cardiovascular imaging, tractography, cell content, vessel wall

## Abstract

In this study we investigated the potential of diffusion tensor imaging (DTI) for providing insight into microstructural changes in arterial tissue by exploring the influence that cell, collagen and elastin content have on fractional anisotropy (FA), mean diffusivity (MD) and tractography. Five ex vivo porcine carotid artery models (n = 6 vessels each) – native, fixed native, collagen degraded, elastin degraded and decellularised – were developed to selectively remove components of arterial microstructure. Intact vessels were imaged at 7 T using a DTI protocol with b = 0 and 800 s/mm^2^ and 10 isotopically distributed directions. FA and MD values were evaluated in the medial layer of vessels and compared across tissue models. FA values measured in native and fixed native vessels were significantly higher (p<0.0001) than those in the elastin degraded and decellularised arteries. Collagen degraded vessels had a significantly higher (p<0.01) FA than elastin degraded and decellularised vessels. Native and fixed vessels had significantly lower (p<0.0001) MD values than elastin degraded, while the MD in decellularised arteries was significantly higher than that in both native (p<0.01) and fixed (p<0.005) tissue. Significantly lower (p<0.005) MD was measured in collagen degraded compared with the elastin degraded model. Tractography results yielded similar helically arranged tracts for native and collagen degraded vessels, whilst elastin degraded and decellularised vessels showed no consistent tracts. FA, MD and tractography were found to be highly sensitive to changes in the microstructural composition of arterial tissue, with cell content being a dominant source of the measured anisotropy in the vessel wall.

## Introduction

Stroke and ischaemic heart disease are the most prevalent forms of cardiovascular disease^1^, while atherosclerosis is widely accepted as the most significant contributor to these burdens^2^. Although numerous mechanisms are associated with the progression of atherosclerosis, changes in vessel microstructure are implicated at the early stages of disease onset^3,4^. As such, imaging markers that are sensitive to changes in arterial tissue microstructure have the potential to provide unique insight into disease onset and progression.

Diffusion tensor imaging (DTI) offers a non-invasive method to probe tissue microstructure, providing quantitative metrics such as mean diffusivity (MD) and fractional anisotropy (FA) which describe the interaction between proton diffusion and the underlying tissue structure. While DTI has predominantly been used to examine white matter, its application in tissue outside the brain has taken significant strides in recent years. To date, a handful of studies have explored the application of DTI to arterial tissue^5,6^ and demonstrated its sensitivity to changes in tissue integrity^7–9^. These studies have laid the groundwork and demonstrated both the feasibility and promise of DTI to effectively investigate underlying tissue microstructure in arterial vessels. However, the effect that microstructural changes have on key diffusion metrics remains unanswered. In a multifaceted microstructure like that of arterial tissue, understanding the impact of the different tissue constituents is critical to inferring any clinically relevant insights from DTI measurements.

Healthy arterial tissue is composed of three main layers: the intimal, medial and adventitial layers. Regardless of location in the body, these layers are predominantly composed of smooth muscle cells (SMCs), collagen and elastin in varying degrees^10^. SMCs attach to elastin fibres and arrange obliquely between concentric lamellae^11^. These SMCs are responsible for the turnover of collagen in the extracellular matrix. This collagen – predominantly type I – is the main load-bearing component of arterial tissue^12^. The close relationship between elastin, SMCs and collagen, together form a continuous helically arranged matrix^10,11^, whose ability to withstand forces both circumferentially and axially allow for the proper mechanical function of healthy arterial tissue. The quantity, quality and arrangement of these components can be disrupted during disease progression and result in significant mechanical shortcomings and failings^4,13,14^.

Fibre tractography has previously been reported to show the helical arrangement of healthy arterial tissue microstructure^5,6^, as well as the disruption to these highly organised tracts when the underlying microstructure is damaged^7^ and the high variability of fibre arrangements in an atherosclerotic plaque^15^. Additionally, Shahid et al.^8^ reported decreased FA values when altering healthy arterial tissue by cutting it and maintaining it open. However, the specificity of FA, MD and tractography to individual microstructural components, such as SMCs, collagen and elastin, remains unclear, therefore, impeding the interpretation of such metrics.

Previous studies on articular cartilage have looked at FA, MD and fibre tractography through the thickness of cartilage^16^ and between anterior and posterior ligaments^17^, where the degree of collagen alignment differs, as well as in damaged cartilage^18,19^, where collagen orientation is disrupted. While these studies look at the global influence of changing tissue structure on DTI metrics, they all show changes in FA, MD and tractography which correlate well, morphologically, to changes in collagen content and arrangement. Fibre tractography has also been shown to be sensitive to the time-dependent orientation of collagen fibres in biodegradable tissue engineered constructs seeded with human-derived vascular cells^20^. Similarly, the orientation of cardiomyocytes^21,22^ has been illustrated by tractography, as well as the differentiation between healthy and diseased cardiac architecture^23–25^. Together, these studies allude to the ability of DTI metrics to be selectively sensitive to specific microstructural components, but this has yet to be conclusively determined in arterial tissue.

The aim of this study is to investigate the potential of DTI to provide specific insight into microstructural changes in arterial tissue by exploring the influence of key components on FA, MD and fibre tractography. This is achieved using ex vivo porcine carotid artery (PCaA) models, developed to selectively remove individual elements of arterial microstructure – SMCs, elastin and collagen. Comparing FA, MD and tractography across these models allows for microstructural insight using DTI metrics. These metrics have the potential to yield novel characterisation of both arterial health and disease progression.

## Methods

### Specimen preparation

Porcine carotid arteries (PCaA) from 6-month-old Large White pigs were excised within three hours of slaughter, cleaned of connective tissue and cryopreserved at a controlled rate of −1°C/min to −80°C in tissue freezing media. Tissue freezing medium was made up of Gibco RPMI 1640 Medium (21875034, BioSciences), sucrose (S0389, Sigma) and the cryoprotectant dimethylsulfoxide (PIER20688, VWR International). Cryoprotectants have been shown to prevent the formation of ice crystals and therefore maintain tissue microstructure during freezing^26,27^. Upon thawing at 37°C, vessels were rinsed in phosphate buffered saline (PBS) (P5493, Sigma) to remove any excess cryoprotectant. Five tissue models were used in this study (n=6 for each): native, fixed native, collagen degraded, elastin degraded and decellularised PCaA. All vessels were cryopreserved prior to treatment and imaged directly after treatment. Table 1 outlines the models and their respective treatments.

**Table 1.** Five different PCaA tissue models and the respective treatments.

| PCaA tissue model | Treatment |
| --- | --- |
| Native | N/A |
| Fixed native | 4% formalin (HT501128, Sigma) fixation for 7 days at 4°C |
| Collagen degraded | 1000 U/ml purified collagenase (CLSPA, Worthington Biochemical Corporation) in MgCl<sub>2</sub> + CaCl<sub>2</sub> supplemented PBS (D8662, Sigma) at 37°C for 28 hours on a rotator |
| Elastin degraded | 10 U/ml purified elastase (ESFF, Worthington Biochemical Corporation) with 0.35 mg/ml trypsin inhibitor (10109886001, Sigma) in Dulbecco's Modified Eagle Medium, high glucose, GlutaMAX (61965026, BioSciences) at 37°C for 3.5 hours |
| Decellularised | 0.1 M sodium hydroxide (S8045, Sigma) perfused through native vessels via a peristaltic pump at 2 Hz for 15 hours, followed by 0.1 M sodium chloride (S3014, Sigma) for 32 hours – all with a pressure of 100 mmHg during perfusion; then treated with 10 µl/ml DNAase (LS006343, Worthington Biochemical Corporation) and 2 µl/ml primicin (Ant-pm-2, InvivoGen) at 37°C for 19 hours<sup>27</sup> |

### MR data acquisition

A small bore (35 mm) horizontal 7T Bruker BioSpec 70/30 USR system (Bruker, Ettlingen Germany) equipped with a receive only 8-channel surface array coil, birdcage design transmit coil, shielded gradients (maximum strength 770 mT/m) and Paravision 6 software was used for all imaging. All vessels were positioned using a custom-made 3D printed holder placed in a 50-ml falcon tube and immersed in fresh PBS prior to imaging at room temperature. A 3D DTI sequence was used with the following parameters: TE/TR: 17.682/1000 ms, image size: 96 × 96 × 60, field of view: 30 × 30 × 18.75 mm, isotropic resolution: 312.5 × 312.5 × 312.5 μm. One b-value of 0 s/mm^2^ was acquired, with a b-value of 800 s/mm^2 5^ subsequently applied in 10 isotopically distributed directions^28^. Equation 1 defines the b-value^29,30^, where γ is the gyromagnetic ratio, G is the gradient amplitude, Δ is the diffusion gradient separation and δ is the diffusion gradient duration.

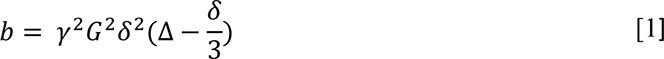

In this study, δ was 3.8 ms, Δ was 8.802 ms and total acquisition time was 17 hours and 36 minutes.

### Image reconstruction and processing

All raw data was denoised^31^ and corrected for Gibbs ringing^32^ in MRtrix3^33^ (www.mrtrix3.org) prior to the mono-exponential tensor model fitting in ExploreDTI^34^. Equation 2 shows the mono-exponential model^29,30^, where *b* is the b-value and *D* is the apparent diffusion coefficient.

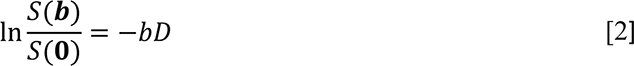

This equation is expanded to incorporate the diffusion tensor, *D*_*ij*_, and the b-matrix, *b*_*ij*_, to produce Equation 3^30^.

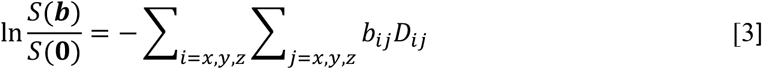

From the tensor, the MD and FA were calculated in ExploreDTI. The MD represents the total diffusion within a voxel, while FA is indicative of the degree of anisotropic diffusion occurring within a voxel on a scale of 0-1^35,36^.

### Regional analysis & tractography

Regions of interest were manually drawn within the media of each vessel using an image created from the mean of the b = 800 s/mm^2^ images. Mean values of FA and MD were calculated within these regions for each vessel. Tractography was similarly performed within ExploreDTI and all parameters used are presented with the corresponding results.

### Histology

For histological processing, samples were fixed in 4% formalin for seven days at 4°C prior to stepwise dehydration in ethanol to xylene. Once dehydrated, all samples were embedded in paraffin wax and sectioned at 8 μm thick slices prior to staining. All stains, their purpose and required imaging are listed in Table 2. All imaging was done using an Olympus BX41 microscope with Ocular V2.0 software. Polarised light microscopy (PLM) uses a polarised filter and two images 90° to each other to maximise the birefringence of collagen for visualisation. Representative histological images are presented.

**Table 2.** Histological stains used in this study, their visualisation and how they are imaged.

| **Stain** | **Visualisation** | **Imaging** |
| --- | --- | --- |
| Alcian blue | Sulphated mucans, such as glycosaminoglycans (GAGs), stain blue | Brightfield |
| Haematoxylin and eosin (H&E) | Haematoxylin stains acidic structures, like cell nuclei, purple-blue and eosin stains basic features, like cytoplasmic filaments, membranes and fibres, pink | Brightfield |
| Picrosirius red | Collagen stains red, while PLM selectively highlights collagen birefringence allowing for visualisation of the orientation | Brightfield + PLM<sup>37</sup> |
| Verhoeff's | Elastin stains black | Brightfield |

### Statistical analysis

Statistical analysis was performed with GraphPad Prism (Version 8). One-way ANOVA with Tukey’s post hoc test for multiple comparisons was used to analyse the variance between groups and determine significance. All numerical and graphical significance is shown as mean ± standard deviation and p<0.05 was considered significant.

## Results

### Tissue model validation

Five tissue models were used in this study to investigate the sensitivity of DTI to the microstructural components of arterial tissue. The fixed native PCaA model is not presented in the histological figures as all PCaA models are fixed prior to histological processing, making it redundant compared to histology of native PCaA. Figure 1 shows H&E, Verhoeff’s and Alcian blue staining for the tissue models. In order to truly understand the influence of each of the tissue components, the selective removal of individual microstructural components was necessary and is confirmed here. H&E verifies cellular content remained in all model tissues, with the exception of decellularised (Figure 1d, top row) – where the complete removal of cells is confirmed. Similarly, the Verhoeff’s elastin stain validates that elastin was removed in the elastin degraded model only (Figure 1c, middle row). Alcian blue (Figure 1, bottom row) shows a variety of GAG concentrations throughout the models. While this hasn’t been investigated in arterial tissue, GAGs have been shown to leach out of tissue when immersed in PBS in order to establish homeostasis^38^.

**Figure 1.**
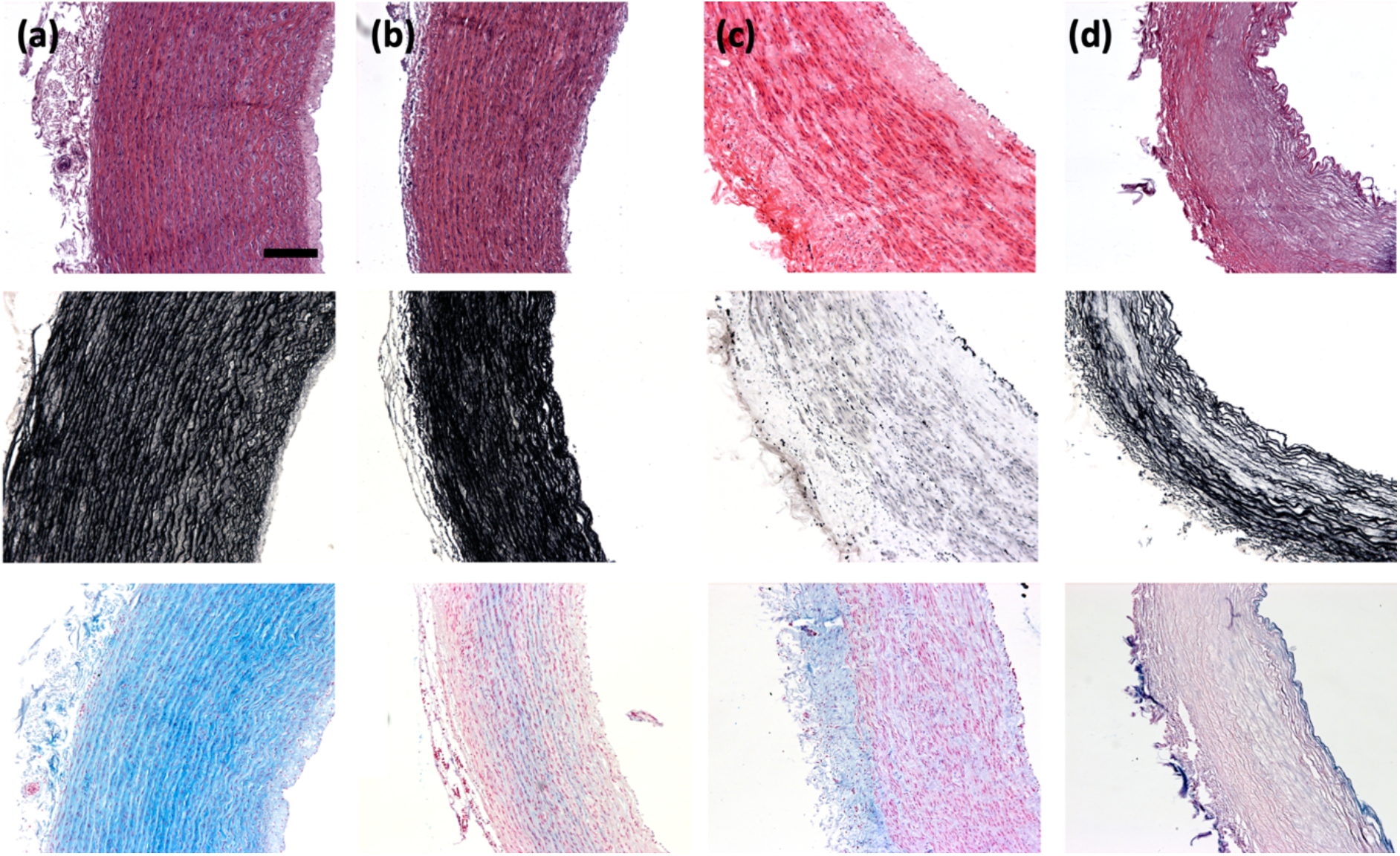
Histological representations of cell, elastin and glycosaminoglycans in (a) native, (b) collagen degraded, (c) elastin degraded and (d) decellularised PCaA. Top to bottom: cell content visible by purple-blue nuclei in H&E, elastin shown in Verhoeff’s elastin stain in black and GAGs stained blue by Alcian blue. All imaged using brightfield microscopy. Scale bar 250 μm.

Figure 2 similarly validates the tissue models, but with respect to collagen content by picrosirius red staining. The top row shows brightfield imaging of the models where collagen is visualised in red. PLM, in the second row, has a specificity for the birefringence of collagen fibres and therefore gives a representation of collagen fibre orientations. Together these confirm that the collagen degraded model (Figure 2b) removed all collagen content. These also confirm that while collagen content was not affected in the other models, neither was the collagen orientation.

**Figure 2.**
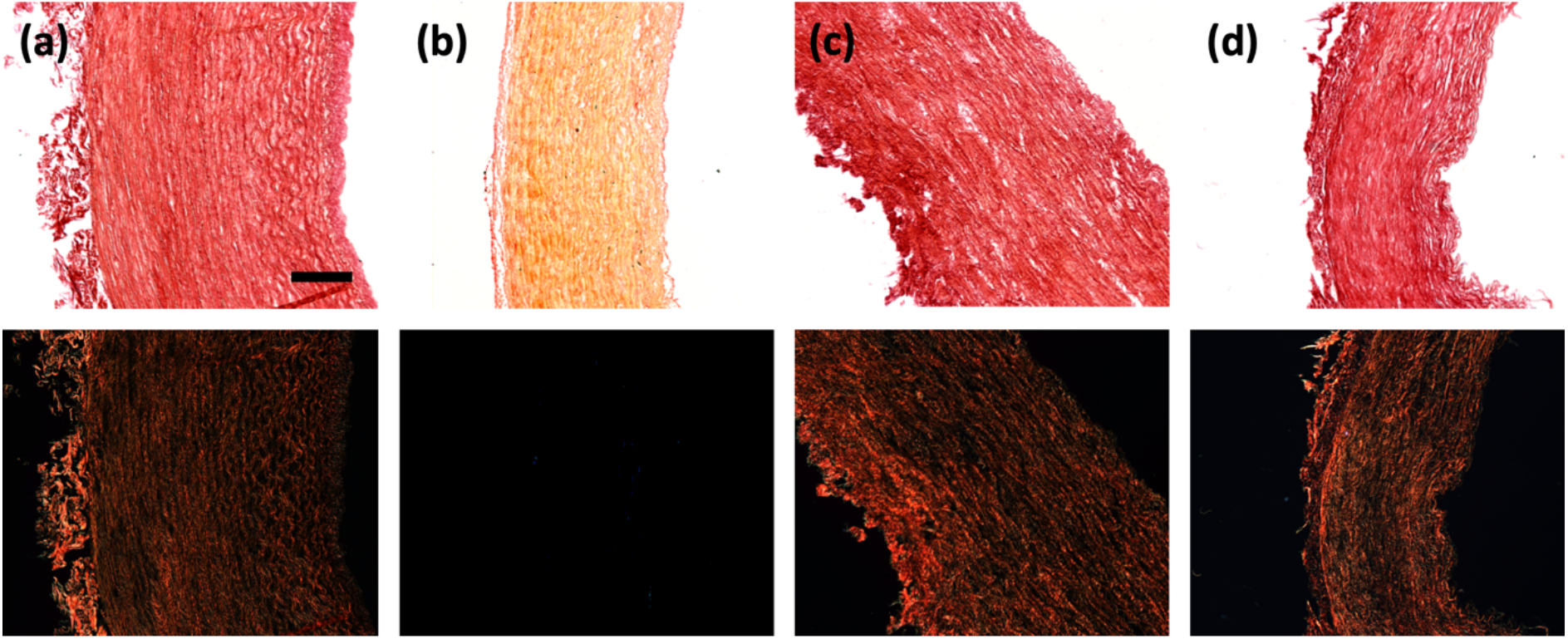
Histological representation of collagen content and orientation in (a) native, (b) collagen degraded, (c) elastin degraded and (d) decellularised PCaA. Brightfield microscopy (top row) shows all tissue stained red, where the PLM on the bottom row has a specificity for the birefringence of collagen. Scale bar 250 μm.

The mean FA measured in the media of each vessel is grouped by tissue model and presented in Figure 3 alongside parametric maps of FA in a representative slice for each model. Visually, the FA maps show stark differences between select tissue models. Native, fixed native and collagen degraded PCaA (Figure 3a, b and c) appear similar. The elastin degraded PCaA (Figure 3d) shows the expansion of the vessels and the apparent loss of tightly bound structure seen in native PCaA. In contrast, both the elastin degraded and decellularised PCaA (Figure 3e) show lower FA ranges. These observations were confirmed by the mean FA measured in the media of each vessel. Native and fixed PCaA demonstrated significantly higher FA than both the elastin degraded and decellularised PCaA (**** p<0.0001). Additionally, the collagen degraded PCaA maintained a significantly higher FA than elastin degraded and decellularised (** p<0.01). No significant differences were seen between native, fixed or collagen degraded PCaA.

**Figure 3.**
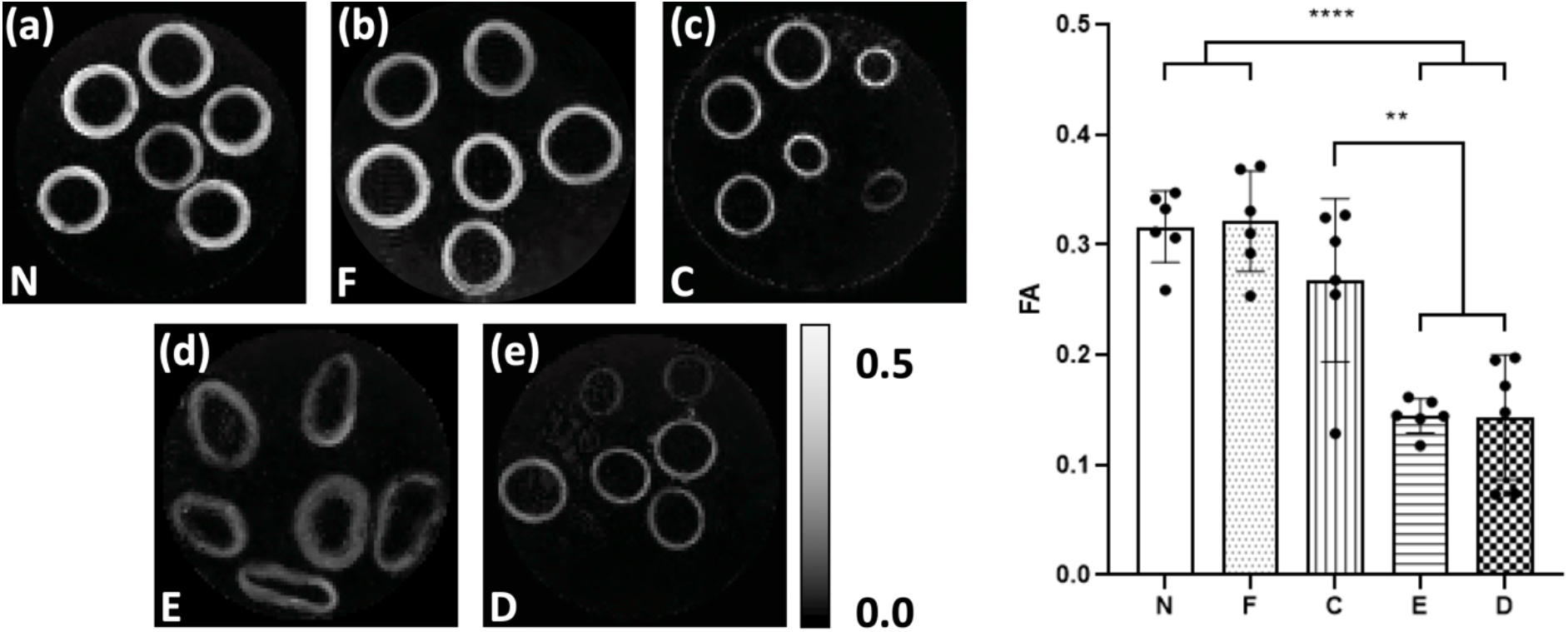
Parametric maps of FA in a representative slice for each of the tissue models. As measured in vessel media, both (a) native (N) and (b) fixed native (F) PCaA showed significantly higher FA than both the (d) elastin degraded (E) and (e) decellularised (D) tissue models. (c) Collagen degraded PCaA also showed a significantly higher FA than both elastin degraded and decellularised PCaA. (** p<0.01, **** p<0.0001)

Parametric maps of MD and regional values of MD extracted from the vessel media for each tissue model are presented in Figure 4. Native and fixed PCaA showed significantly lower MD values than the elastin degraded model (**** p<0.0001), which can be seen visually in the MD maps (Figure 4a, b). Decellularised PCaA (Figure 4e) had a significantly higher overall diffusion than both native (** p<0.01) and fixed tissue (*** p<0.005). Fixed PCaA had a lower mean MD than native PCaA, however no significant difference was found. The collagen degraded PCaA showed a significantly lower MD than the elastin degraded model (*** p<0.005), which is evidenced in the MD maps as well (Figure 4c, d).

**Figure 4.**
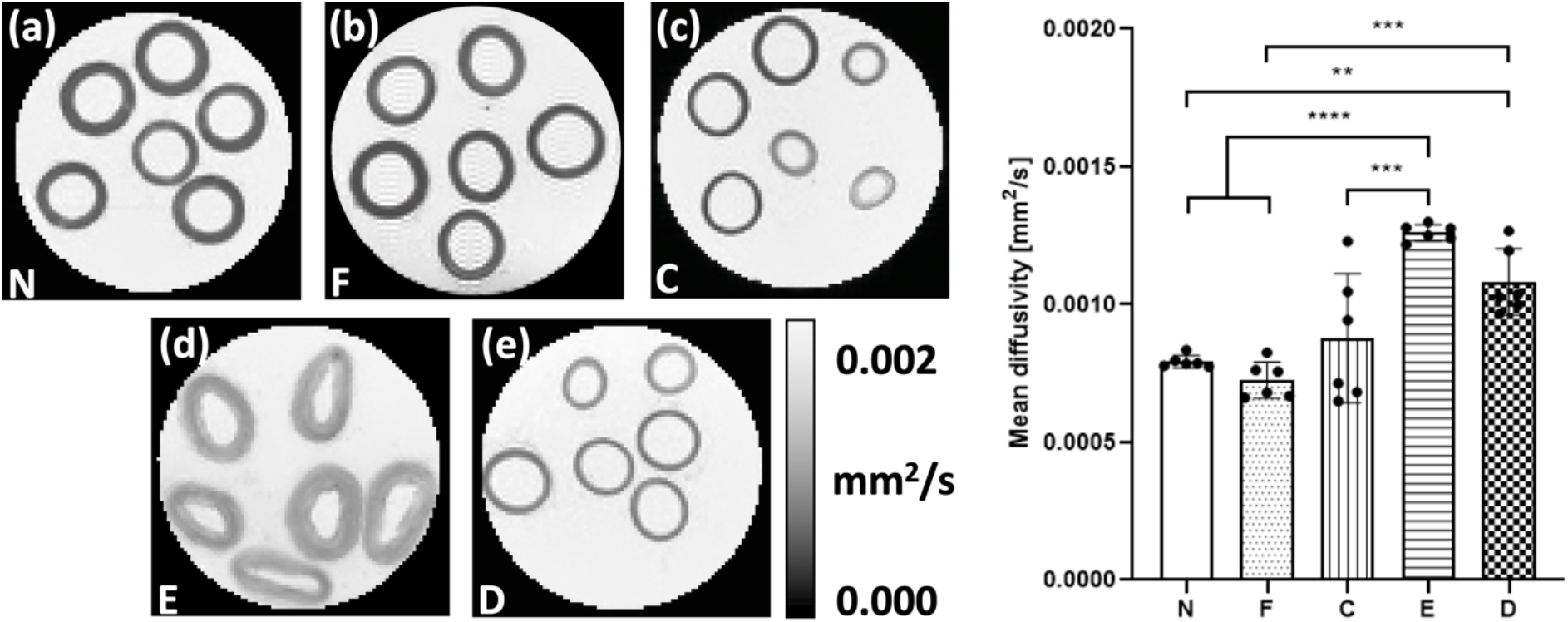
Parametric maps of MD in a representative slice for each of the tissue models. As measured in vessel media, both (a) native (N) and (b) fixed native (F) PCaA showed a significantly lower MD than both the (d) elastin degraded (E) and (e) decellularised (D) PCaA. The elastin degraded PCaA had a significantly higher MD than the (c) collagen degraded PCaA. (** p<0.01, *** p<0.005, **** p<0.0001)

Tractography was performed to visualise the diffusion pathways within the tissue models. Keeping tractography parameters constant, Figure 5 demonstrates the varying results obtained across tissue models. Fixed PCaA was not included due to the similarity in tract quality with native PCaA. Native and collagen degraded PCaA (Figure 5a, b) both illustrate a coherent and helical arrangement of tracts which align with the known helical arterial tissue structure. Using the same parameters, the elastin degraded and decellularised PCaA models (Figure 5c, d) show no helically arranged tracts.

**Figure 5.**
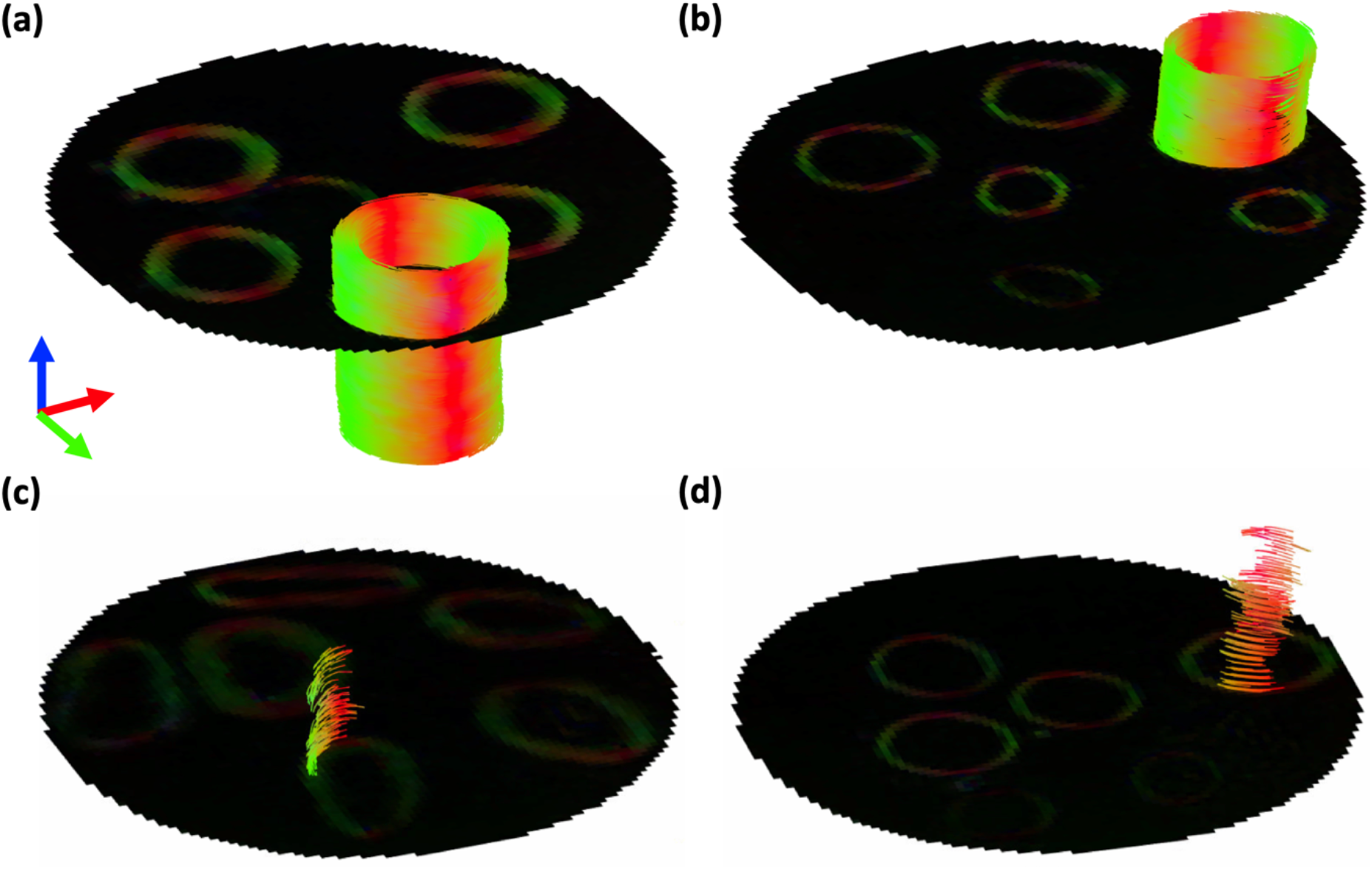
Tractography of (a) native, (b) collagen degraded, (c) elastin degraded and (d) decellularised PCaA. All models were modelled with the following parameters: seed point resolution: 0.3125 mm × 0.3125 mm × 0.3125 mm, FA threshold: 0.2, FA tracking threshold: 0.2 – 1, tract length: 0.5 – 5.0 mm, angular threshold: 30° and step size of 0.3125 mm.

Comparing native and collagen degraded PCaA models, Figure 6 shows tractography for a representative vessel of each model alongside H&E and PLM histology. A cross-sectional view of native PCaA shows the circumferentially aligned cell and collagen fibre content (Figure 6b, c). The histologically verified orientation of both cells and collagen coincide well with the arrangement of the tracts of native PCaA (Figure 6a). The collagen degraded PCaA resulted in similar tracts (Figure 6d), despite the lack of any collagen (Figure 6f), while the circumferentially aligned cell content is still visible (Figure 6e).

**Figure 6.**
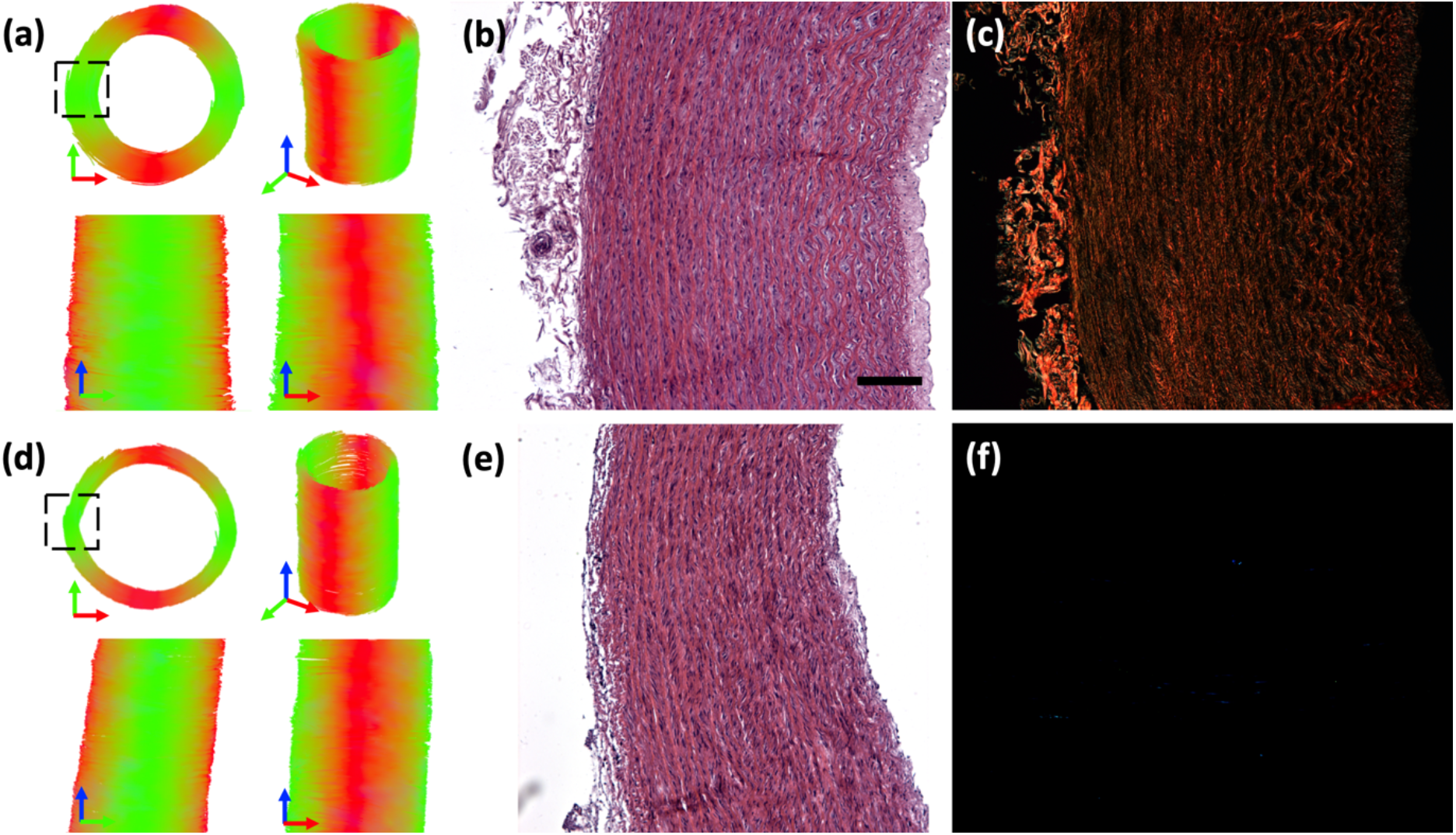
Tractography of representative (a) native and (d) collagen degraded PCaA. Both models were obtained with the following parameters: seed point resolution: 0.3125 mm × 0.3125 mm × 0.3125 mm, FA threshold: 0.2, FA tracking threshold: 0.2 – 1, tract length: 0.5 – 5.0 mm, angular threshold: 30° and step size of 0.3125 mm. Representative cross-sectional histology shows (b, e) cellular arrangement by H&E for native and collagen degraded PCaA, respectively. (c) PLM shows the similar orientation of collagen in native PCaA, and lack thereof in (f) collagen degraded PCaA. Scale bar is 250 μm.

## Discussion

In the present study we investigated the sensitivity of DTI derived FA, MD and tractography to changes in arterial tissue microstructure. By selectively removing SMCs, elastin and collagen we explored how each component plays a part in the typically anisotropic diffusion profile of arterial tissue^8^ (Figure 3). Results from decellularised arterial tissue demonstrate that the main contributor to this anisotropic diffusion in arterial tissue is the presence of cell content. While previous studies highlight the role of collagen fibres in diffusion derived metrics^8,20,39^, here, we evidence the co-dependency of collagen and cell content and characterise their influence on FA, MD and tractography. With the removal of collagen there is no significant change in FA, MD or tractography compared to native arterial tissue. However, the loss of cellular content results in a purely isotropic diffusion, seen by increased MD and decreased FA, despite the presence of collagen fibres as confirmed by histology. This becomes even more evident in the tractography results – where the decellularised vessels yield no helically arranged tracts (Figure 5). This result highlights the significance of cellular content and corroborates findings in a previous ex vivo DTI study on cell migration in brain tumours^40^, where the authors saw a decrease in FA and increase in MD when cells migrated out of a region of interest.

Vascular SMCs on average are 200 μm long and 5 μm in diameter^41^, while a single collagen fibril diameter is 80 ± 11 nm and a fibre bundle is approximately 5.1 ± 6.1 μm^42^. SMCs are responsible for the turnover of the extracellular matrix, including collagen, and therefore their orientation aligns with that of collagen. Together these components form the helical matrix of arterial tissue which has been well documented^5,6^. As our samples were imaged at room temperature, the MD for PBS was found to be 0.00185 ± 0.00001 mm^2^/s. Using the 3D root-mean square equation^43^ 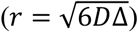 and the gradient interval time used in this study (Δ = 8.802 ms), a diffusing water molecule could travel approximately 9.8 μm, suggesting sensitivity to diffusion at the scale of both SMCs and collagen.

In the absence of obstructing boundaries, protons diffuse freely in all directions. The presence of boundaries, such as SMCs or collagen fibres, cause either restricted or hindered diffusion. Intracellular diffusion is typically regarded as restricted diffusion as the molecules are trapped within the cell membrane and have limited area to diffuse within^44^. The cell membrane is composed of a phospholipid bilayer which is selectively permeable and due to the polar nature of water molecules, they therefore diffuse at a slow rate^45^. Hindered diffusion arises when the diffusion of water molecules is impeded by boundaries, which reduce their net displacement. Generally, extracellular diffusion in biological tissue is characterised as hindered diffusion^44,46^. Within the scale of diffusion presented here, we see the combined effects of both hindered and restricted diffusion^47^ and changes in their joint profile as components are removed.

The removal of SMCs from arterial tissue resulted in a drastic decrease in FA, as can be seen in the decellularised tissue model (Figure 3). The high FA measured in the collagen degraded vessels suggests that SMCs, and the intracellular restricted diffusion attributed to them, are the main contributors to this anisotropic diffusion– more so than the hindered diffusion from interactions with collagen fibres. Removal of elastin from the artery resulted in the most isotropic diffusion of all models, seen by low FA and the highest MD. While the quantity of cells did not change in the elastin degraded model, the removal of elastin resulted in a less compact and organised cell alignment compared to native arterial tissue – this can be seen histologically by H&E (Figure 1a, c). SMCs attach to the concentric elastin lamellae and are embedded between collagen fibres^11^. The removal of elastin resulted in an expansion in the extracellular space, which enhances proton mobility and, therefore, increases the MD^48^. This structural change can be seen plainly in the FA (Figure 3d) and MD (Figure 4d) maps. The MD of the elastin degraded model is higher than that of the decellularised model suggesting that not only does the less densely packed cell content and arrangement affect the DTI metrics compared to native, but the decrease in hindered diffusion from the removed elastin molecules also plays a significant role in this isotropic response.

It is worth noting that all the tissue models displayed a decrease in GAG content (Figure 1, bottom row). A study by Bartholomew and Anderson^49^ demonstrated that proteoglycans, the proteins GAGs attach to, coat collagen type III in the media, which in turn coats the elastin fibres. This suggests it is not possible to avoid the disruption and depletion of GAGs in arterial tissue when removing collagen or elastin. In cartilage, Xia, et al.^48^ illustrated that the MD has no direct correlation to GAG content but instead they proposed that the space left from degraded and removed macromolecules allows for increased diffusivity – which has since been demonstrated^50^. To the author’s knowledge, no studies exist examining the influence that GAGs have on diffusion in arterial tissue; however, while proteoglycans show no preferred orientation and therefore shouldn’t influence FA, as previously suggested, their removal could increase MD as a result of the open extracellular space left behind upon their removal.

The main constituents of native PCaA are SMCs, elastin and collagen. When fixing native arterial tissue, these constituents were unaffected and there was no effect on the FA, MD or tractography. While multiple studies have reported increased MD in cardiac tissue when fixed^9,51,52^, the length of fixation, concentration of fixative and time after fixation in cardiac tissue have resulted in considerable variation in measured FA and MD^53,54^. One study on fixed tissue observed an initial decrease in the MD followed by an increase after 15 days^53^, whilst it has also been shown that increased exposure to fixatives can cause cell membrane degradation by the depletion of lipid membranes through carbon double bond reactions^55^. Our results showed no significant difference between fresh and fixed tissue with respect to the FA, MD and tractography. The length of our fixation protocol, 7 days, is likely short enough to avoid any membrane degradation and therefore had no significant effect on diffusivity.

Previous studies in arterial tissue have looked at the structure of native arterial tissue as well as how storage and preparation for imaging effect the diffusion profiles. While the fibre angles have not been quantified in the present study, the helically aligned tractography of native and collagen degraded tissue corroborate previous studies highlighting the helical arrangement of SMCs and collagen^5,6^. The results from the tractography analysis provide visualisation of the significance of cell content on the diffusion profile. It is demonstrated in this study that, within arterial tissue, tractography is far more indicative of cellular orientation over collagen fibre arrangement. Histological analysis demonstrates that SMCs and collagen follow the same circumferential arrangement in native arterial tissue (Figure 6a) and tractography yields good visualisation of that structure. However, for the collagen degraded vessels (Figure 6d) the tractography is a representation of cellular alignment alone.

Microstructural changes in arterial tissue can have significant implications for the mechanical functionality of the tissue^56^, as well as often being a precursor to disease progression^4^. On this basis and considering the characterisation of arterial microstructure presented in this study, DTI has the potential to provide unique biomarkers for the integrity of arterial tissue. Atherosclerosis is a chronic immunoinflammatory, fibroproliferative disease which starts with the adhesion of low-density lipoproteins to the intimal layer of the arterial wall^3^. After this initial step, macrophages and foams cells rapidly accumulate at the intimal layer and migrate into the intimal-medial layer boundary resulting in a continually changing microstructure as disease progresses. Morphologically, the first signs of an atherosclerotic plaque are the thickening of the intima, followed by the well-known formation of a lipid core and fibrous cap^57,58^. Microstructurally, these different regions have altered quantities and arrangements of SMCs, collagen and elastin. The thickened intima typically shows a decrease in SMC content^57^, the lipid core highlights the displacement of SMCs by foam cells^58^, and the fibrous cap, which covers the lipid core^59^, has been shown to have quite variable SMC content depending on location^60^. The demonstrated sensitivity of DTI to SMC content in arterial tissue in this study suggests that it may be an ideal metric to identify such early indicators of disease driven microstructural changes. Additionally, other cardiovascular diseases – such as aneurysms – have shown significant fragmentation of the elastic lamellae which can cause catastrophic failure of the arterial wall^61^. Changes in the key microstructural components of arterial vessels can lead to significant mechanical failings^56^ and identifying these changes using imaging biomarkers, offers potential insight into the mechanical integrity of the arterial wall in atherosclerotic^13,62,63^ and/or aneurysmal^61^ tissue.

Few studies have looked at the implementation of diffusion imaging in vivo for carotid artery and atherosclerotic plaque visualisation^64–67^ as there are many elements which make clinical translation challenging. The high resolution, lack of physiological motion^68^ and extended scan time in this study allowed for a detailed look at the vessel microstructure, which would be necessary for investigating regions of atherosclerotic plaques. Despite the idealised ex vivo experimental set up, the work presented here highlights the promise for DTI metrics to yield valuable insight into arterial microstructure which could ultimately provide novel insight into diseased tissue morphologies. For example, recent in vivo studies have used quantitative susceptibility mapping to investigate gross morphological features^69–71^ and inflammation^72^ in atherosclerotic plaques, but this approach also has potential to provide markers of tissue microstructure and integrity^73^. Ideally, a combination of methods which allow for the full characterisation of the microstructure within the vessel wall would provide the insight needed to better inform the risk of atherosclerotic plaque rupture. This study establishes the influence of key microstructural components on diffusion metrics in arterial tissue and highlights the potential of DTI for identifying disease driven changes in arterial microstructure.

## Abbreviations

DTI: diffusion tensor imaging
FA: fractional anisotropy
MD: mean diffusivity
SMCs: smooth muscle cells
PCaA: porcine carotid arteries
PBS: phosphate buffered saline
PLM: polarised light microscopy
H&E: Haematoxylin and eosin
GAGs: glycosaminoglycans

## Acknowledgements

This research was funded by the European Research Council (ERC) under the European Union’s Horizon 2020 research innovation programme (Grant Agreement No. 637674).

